# Suppressing Bone Resorption and Promoting Mineralization with Tetracycline Derivatives

**DOI:** 10.64898/2026.02.24.707630

**Authors:** Saeka Shimochi, Katja Hrovat, Uruj Sarwer, Leire Bergara-Muguruza, Ari Alho, Maarit Laine, Akihisa Otaka, Yasuhiko Iwasaki, Ilkka Paatero, Ulvi K. Gursoy, Keijo Mäkelä, Eriika Savontaus, Jukka Salonen, Miho Nakamura

**Author notes:** Funding: Sigrid Jusélius Foundation (#230131), the Japan Society for the Promotion of Science (#23K08670), the Murata Science Foundation, the Turku Collegium for Science, Medicine and Technology.

## Abstract

Osteoporosis is a progressive skeletal disorder characterized by decreased bone mass and an increased risk of fracture. Current treatments are limited by adverse effects and poor long-term compliance, necessitating alternative therapeutic approaches. Tetracycline (TC) derivatives, which are traditionally used as antibiotics, have shown promise in modulating bone remodeling. In this study, the effects of TC and three TC derivatives—oxytetracycline (OC), doxycycline (DC), and minocycline (MC)—on osteoclast and osteoblast activities were investigated using in vitro human cell models and in vivo zebrafish assays. All TC derivatives inhibited osteoclast differentiation and bone resorption, as shown by reductions in the number of TRAP-positive cells, resorption pit volume, and matrix metalloproteinase (MMP)-2/MMP-9 secretion. DC demonstrated the most potent inhibitory effects across all concentrations. Low to moderate concentrations of OC, DC, and MC promoted osteoblast proliferation and mineralization, whereas high doses inhibited these processes. Confocal imaging confirmed the accumulation of TC derivatives in mineralized bone nodules. Zebrafish studies revealed dose-dependent suppression of craniofacial bone development at higher concentrations. These findings highlight the dose dependent, dual effects of TC derivatives on bone cells (osteoblasts and osteoclasts) and underscore the potential of these agents as dual-function therapies for osteoporosis.

## 1. Introduction

Osteoporosis is a chronic and progressive metabolic bone disease characterized by decreased bone mass and deterioration of the bone microarchitecture, leading to increased bone fragility and fracture risk. Globally, osteoporosis is a growing health concern, particularly in aging populations. Epidemiological data indicate that one in three women and one in five men will experience osteoporotic fractures in their lifetime ^[1]^. As life expectancy increases, the prevalence and socioeconomic burden of osteoporosis are expected to increase significantly, underscoring the urgent need for safe, effective, and affordable therapies. Current osteoporosis treatments are predominantly categorized into antiresorptive and anabolic agents. Bisphosphonates, nuclear factor kappa-B ligand (RANKL) antibodies, selective estrogen receptor modulators, and calcitonin are commonly used as antiresorptive drugs, while anabolic options include parathyroid hormone analogs and monoclonal sclerostin antibodies. Despite their proven efficacy, these treatments have considerable limitations.

Long-term bisphosphonate use is linked to complications such as atypical femoral fractures, osteonecrosis of the jaw, and a risk of osteosarcoma development ^[2–5]^. The long-term use of bisphosphonates for more than 6 years is associated with higher fracture rates ^[2]^ and a high risk of osteonecrosis ^[3]^, atypical fracture ^[4]^ and osteosarcoma ^[5]^. To minimize these risks, drug holidays for 1–2 years are needed. Furthermore, RANKL antibodies, while effective at increasing bone mineral density, carry a heightened risk of infections because of their immunomodulatory effects ^[6]^. These adverse effects often necessitate treatment holidays and can limit long-term compliance and efficacy. Although the monoclonal RANKL antibody has advantages over bisphosphonates in promoting bone density, it carries a risk of infection because the target is part of the immune system; additionally, this treatment is associated with the development of osteonecrosis of the jaw ^[6]^. Most osteoporosis treatments are limited by side effects, serious risks and high costs. Monoclonal RANKL and sclerostin antibodies were approved for the market in 2010 and 2019, respectively; however, there is still a great need for effective osteoporosis treatments.

Given the need for alternative therapies with improved safety profiles and reduced costs, the potential for repurposing tetracyclines (TCs) as modulators of bone remodeling has attracted increased attention. Initially introduced in the 1950s as broad-spectrum antibiotics, TC and TC derivatives such as oxytetracycline (OC), doxycycline (DC), and minocycline (MC) have since been shown to exert nonantibiotic biological effects ^[7]^. Recently, DC, a semisynthetic TC compound, was reported to exhibit anticancer ^[8]^ and bone repair effects in vivo ^[9]^. At the cellular level, the soluble form of chemically modified TC induces osteoclast apoptosis in vitro ^[10]^ by suppressing matrix metalloproteinase (MMP)-mediated histone H3 cleavage ^[11]^. More recently, a role of these compounds in bone metabolism has been proposed, suggesting that they are potential candidates for osteoporosis therapy. Therefore, TCs could be strong candidates for preventing osteoporosis through the inhibition of osteoclast activity.

Currently, TCs are divided into three groups based on their pharmacokinetic and antibacterial properties ^[12]^. Compounds in group 1, including TC and OC, are characterized by reduced absorption, and these agents are less lipophilic than new TC derivatives. Compounds in group 2, including DC and MC, are essentially completely absorbed and more lipophilic than group 1 TC derivatives. Compounds in group 3 show activity in vitro against bacteria that are resistant to other TC derivatives. The biological activity of these compounds may be enhanced by chemically modifying the upper peripheral zone, particularly at positions C7 through C9 of the D ring ^[13,14]^. This has been accomplished with semisynthetic TC compounds such as DC and MC, in which the upper functional groups were synthetically modified to enhance biological targets ^[13,14]^. However, to retain their antibiotic effects, they must possess a C4 dimethylamino group. New 4-demethylamino TCs and chemically modified TCs have been shown to exert stronger nonantibiotic effects than naturally occurring and semisynthetic TCs and to lack antimicrobial activity ^[13–15]^.

Bone is a dynamic tissue that is continually formed by osteoblasts and resorbed by osteoclasts ^[16]^. The bone remodeling cycle is affected by many factors, including mechanical stress ^[17]^, metabolic factors ^[18]^, endocrine changes, and pharmaceuticals ^[16]^. Considering the importance of a balance between osteoblasts and osteoclasts, the effects of TC derivatives on both types of cells are of great interest. Thus, the aim of the present study was to assess the effects of TC derivatives on the activity of osteoclasts and osteoblasts.

## 2. Results

### 2.1. Effects on osteoclastogenesis

To compare the effects of TC derivatives on osteoclast formation and resorptive activity, human osteoclasts were cultured on bone slices with different types of TC derivatives at several concentrations. TRAP staining revealed that a variable number of bone marrow cells differentiated into osteoclasts (**Figure 1a**). There were TRAP-positive cells with single nucleus, suggesting incomplete differentiation into mature osteoclasts. In this study, TRAP-positive cells with more than 3 nuclei were counted as osteoclasts. The number of multinuclear TRAP-positive cells was significantly greater on bone slices not treated with any TC derivatives (control) and those treated with a low concentration of TC (5 μg/ml). (**Figure 1b**) Dose-dependent decreases in the number of multinuclear TRAP-positive cells were observed in osteoclasts cultured with TC, OC, and MC. DC inhibited the formation of multinuclear TRAP-positive cells at all concentrations (5, 10 and 20 μg/ml).

**Figure 1.**
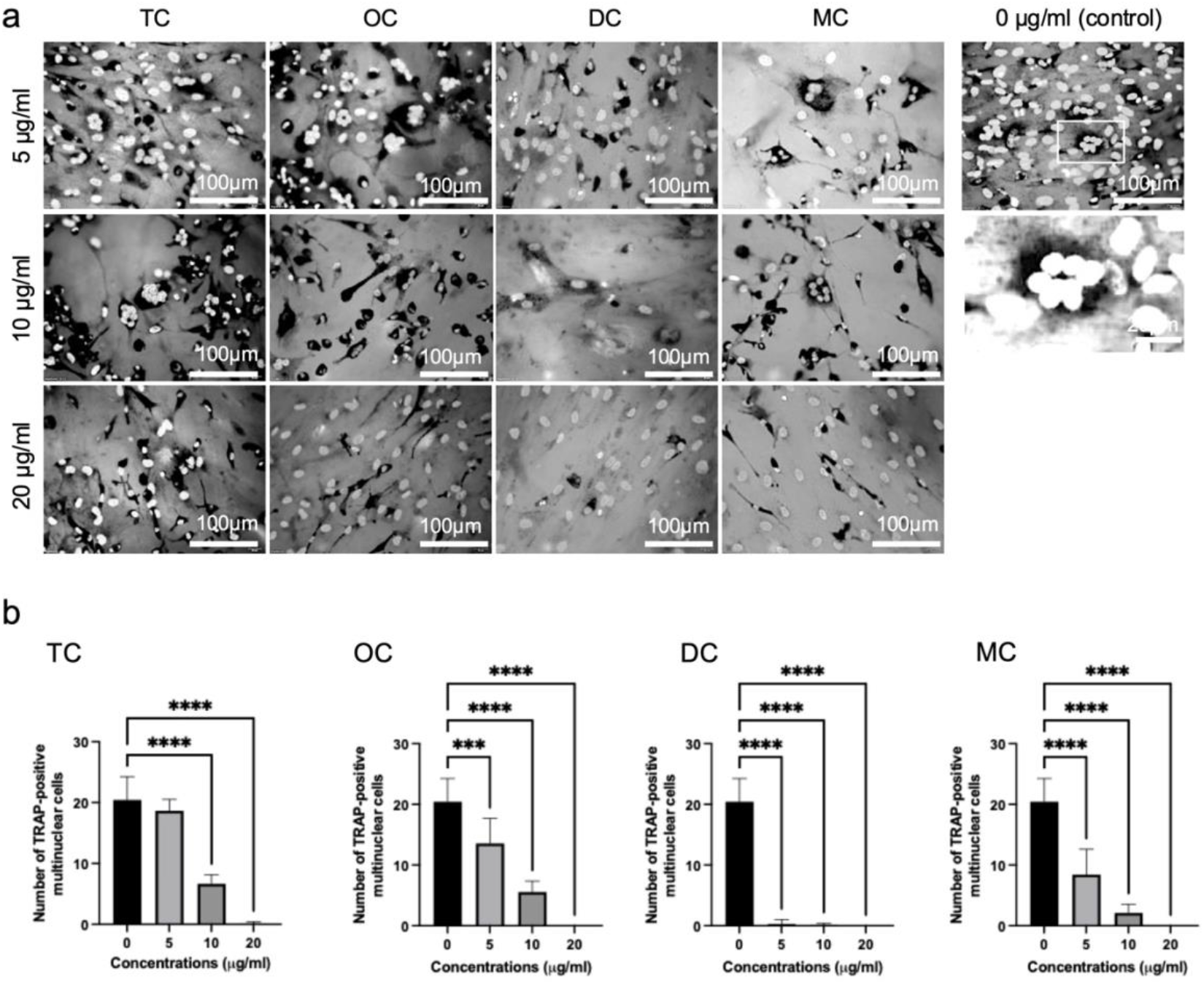
Effects of TC derivatives on osteoclast formation in human bone marrow– derived cultures. (a) Representative TRAP–stained images of human osteoclast cultures grown on bone slices and treated with different TC derivatives at various concentrations. TRAP-positive cells with a single nucleus indicate incomplete differentiation, whereas multinucleated TRAP-positive cells represent mature osteoclasts. (b) Quantification of multinucleated TRAP-positive osteoclasts (cells containing more than three nuclei) under each treatment condition. The control group and cultures treated with low-dose TC (5 μg/mL) exhibited significantly higher numbers of osteoclasts. In contrast, TC, OC, and MC induced dose-dependent reductions in osteoclast formation, while DC markedly inhibited multinucleated osteoclast formation at all tested concentrations (5, 10, and 20 μg/mL). Data are shown as the mean ± SD (n=9). *** *p* < 0.0005, \*\*\*\**p* < 0.0001 compared with the control. n.s.=not significant *p* > 0.05.

Osteoclast resorption pits on bone slices were analyzed by laser microscopy (**Figure 2a**). The most distinct resorption pits formed on bone slices not treated with any TC derivatives (control) and those treated with a low concentration of TC (5 μg/ml) or OC (5 μg/ml). Dose-dependent inhibition of resorption was observed in osteoclasts cultured with TC, OC and MC (**Figure 2b**). The volume of the resorption pits formed on bone slices by osteoclasts treated with a low concentration of TC, OC and MC (5 μg/ml) was approximately 0.7-fold smaller than that in the control group, and a moderate concentration of TC, OC and MC (10 μg/ml) resulted in pits whose volume was 0.1-fold smaller than that in the control group. DC inhibited osteoclast resorption at low, moderate and high concentrations (5, 10 and 20 μg/ml, respectively). Compared with those in the control group, the volume of resorption pits decreased approximately 0.1–0.2-fold after treatment with 10 μg/ml TC, OC and MC. There were very few resorption pits on the bone slices treated with 20 μg/ml TC, OC, DC and MC.

**Figure 2.**
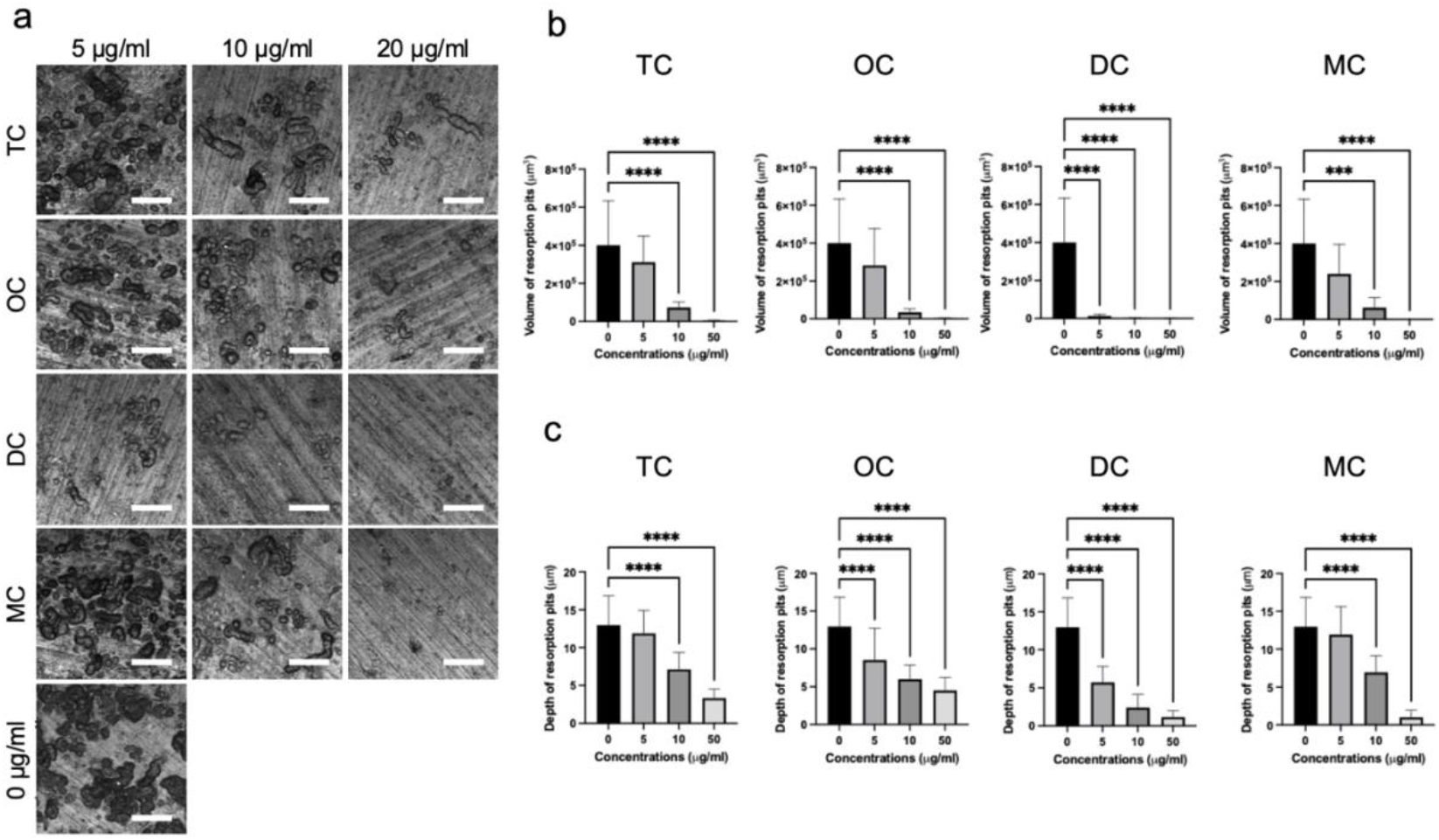
Effects of TC derivatives on osteoclast resorptive activity on bone slices. (a) Representative laser microscopy images of osteoclast-mediated resorption pits formed on bone slices following treatment with different TC derivatives at varying concentrations. Prominent resorption pits were observed in the control group and in cultures treated with low concentrations of TC and OC (5 μg/mL). (b) Quantitative analysis of resorption pit volume under each treatment condition. Osteoclast resorption activity decreased in a dose-dependent manner in response to TC, OC, and MC. Low concentrations (5 μg/mL) of TC, OC, and MC reduced pit volume to approximately 70% of the control, while moderate concentrations (10 μg/mL) resulted in a marked reduction to approximately 10% of control levels. DC significantly inhibited resorption at all tested concentrations (5, 10, and 20 μg/mL). At the highest concentration (20 μg/mL), very few resorption pits were observed across all TC derivative treatments. Data are shown as the mean ± SD (n=9 for volume measurements and n=45 for depth measurements). *** *p* < 0.002, \*\*\*\**p* < 0.0001 compared with the control.

The depth of resorption pits formed by osteoclasts was significantly greater on bone slices not treated with any TC derivatives (control) and those treated with a low concentration of TC (5 μg/ml) and MC (5 μg/ml) (**Figure 2c**). The depth of resorption pits decreased approximately 0.5-fold after treatment with 10 μg/ml TC, OC and MC and approximately 0.2-fold after treatment with 10 μg/ml DC compared with that in the control group. The depth of resorption pits decreased approximately 0.3-fold after treatment with 20 μg/ml TC or OC and approximately 0.1-fold after treatment with 20 μg/ml DC and MC compared with that in the control group.

The concentrations of MMP-2 and MMP-9 in cell lysates were assessed after osteoclast culture on bone slices with different types of TC derivatives for 14 days (**Figure 3a and 3b, respectively**). The secretion of MMP-2 was inhibited in osteoclasts cultured with higher concentrations of TC (10 or 20 μg/ml), OC (10 or 20 μg/ml), and MC (20 μg/ml). DC inhibited the secretion of MMP-2 at all concentrations (5, 10 and 20 μg/ml). The secretion of MMP-9 was inhibited in osteoclasts cultured with all the tested TC derivatives. All types of TC derivatives, even at relatively low concentrations, strongly inhibited MMP-9 secretion from osteoclasts.

**Figure 3.**
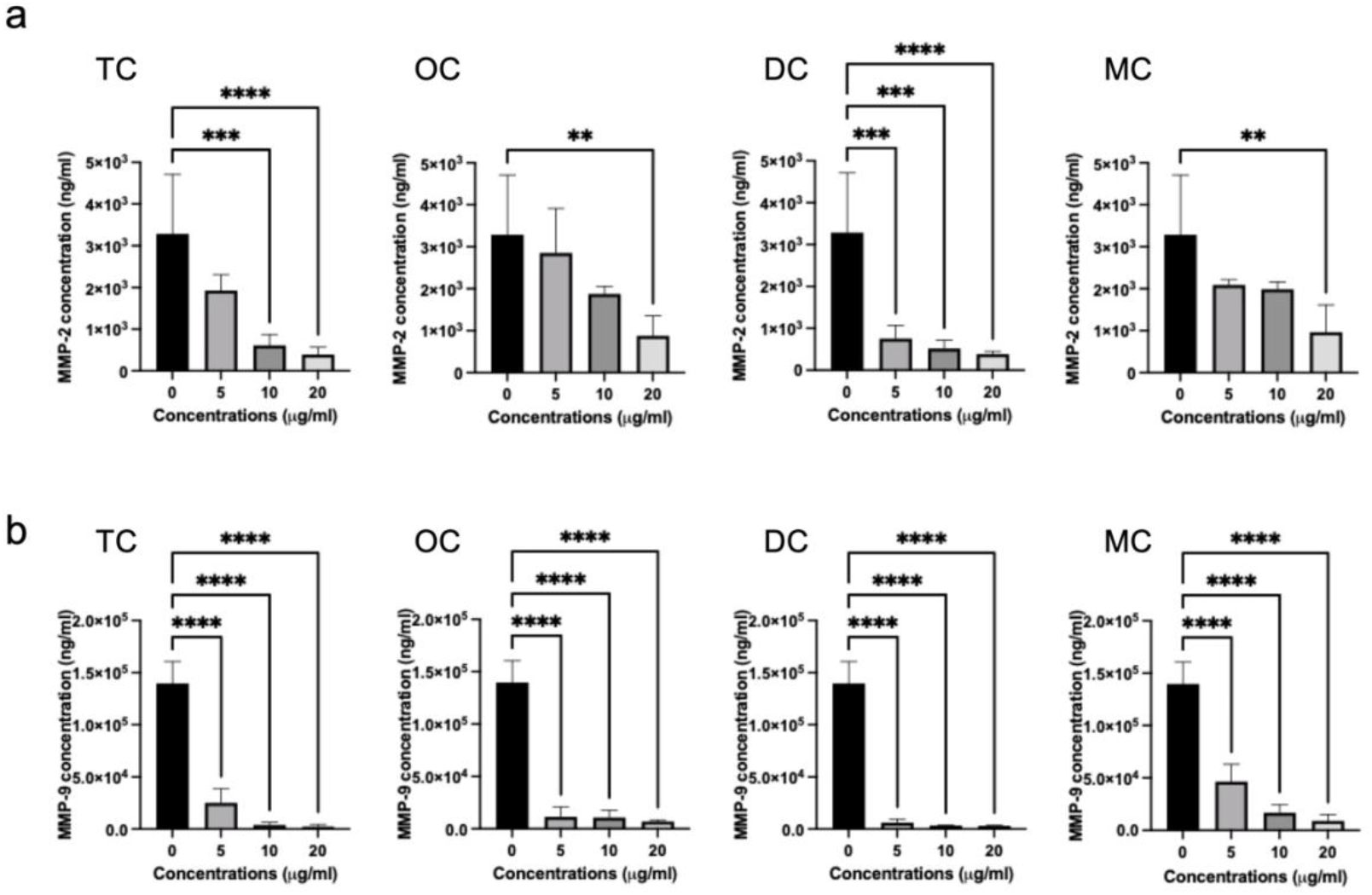
Effects of TC derivatives on MMP secretion in osteoclast cultures. (a) Quantification of MMP-2 levels in cell lysates following 14 days of osteoclast culture on bone slices in the presence of different TC derivatives at varying concentrations. Secretion of MMP-2 was significantly reduced in cultures treated with higher concentrations of TC (10 and 20 μg/mL), oxytetracycline (OC; 10 and 20 μg/mL), and minocycline (MC; 20 μg/mL), while doxycycline (DC) markedly inhibited MMP-2 secretion at all tested concentrations (5, 10, and 20 μg/mL). (b) Quantification of MMP-9 levels in cell lysates under the same experimental conditions. All tested TC derivatives significantly suppressed MMP-9 secretion, with strong inhibitory effects observed even at low concentrations. Data are shown as the mean ± SD (n=5). \*\**p* < 0.005, *** *p* < 0.002, \*\*\*\**p* < 0.0001 compared with the control.

### 2.2. Effects on osteoblast proliferation and differentiation

To compare the effects of TC derivatives on osteoblast proliferation, cell numbers were measured after cell culture for 1 d, 3 d, and 7 d. The cells adhered to and proliferated in wells with the different types of TC derivatives at several concentrations (**Figure 4**). There were no significant differences in the number of cells among all the tested TC derivatives at 1 d after seeding. The number of cells gradually increased after they attached to the well. The cell numbers were greater after treatment with a low and moderate concentration (5 and 10 μg/ml) of TC, DC and MC at 3 d after seeding, whereas the cell numbers were the same after treatment with a high concentration of all the tested TC derivatives (20 μg/ml). During the late phase of proliferation at 7 d, the cell numbers were greater in the TC group (5 and 10 μg/ml); however, the cell number was not different in the other TC derivative groups. OC did not affect cell proliferation at any time point.

**Figure 4.**
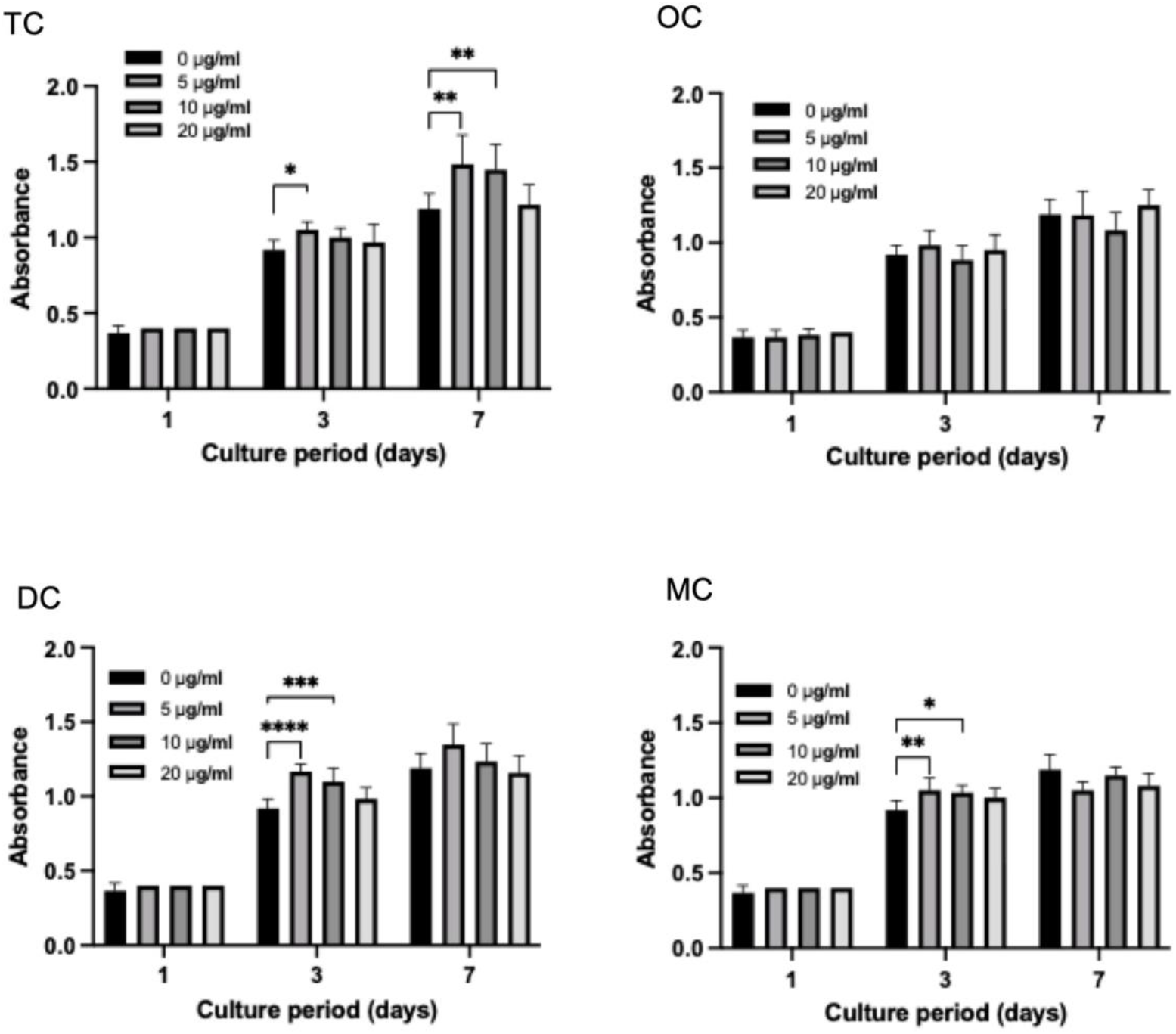
Effects of TC derivatives on osteoblast proliferation. Osteoblast cell numbers were quantified after 1, 3, and 7 days of culture in the presence of different TC derivatives at varying concentrations. Cells adhered and proliferated under all treatment conditions. No significant differences in cell numbers were observed among groups at 1 day after seeding. At 3 days, cultures treated with low and moderate concentrations (5 and 10 μg/mL) of TC, DC, and MC. exhibited increased cell numbers compared with the control, whereas high concentrations (20 μg/mL) of all TC derivatives showed no effect on proliferation. At 7 days, enhanced cell proliferation was maintained in the TC-treated groups at 5 and 10 μg/mL, while no significant differences were observed in the other derivative-treated groups. Oxytetracycline (OC) did not affect osteoblast proliferation at any time point. Data are shown as the mean ± SD (n=10). \**p* < 0.02, \*\**p* < 0.005, *** *p* < 0.002, \*\*\*\**p* < 0.0001 compared with the control.

To compare the effects of TC derivatives on osteoblast differentiation, osteoblasts were cultured with different types of TC derivatives at several concentrations. The cells not treated with any TC derivatives (control) began mineralization at 4 weeks and then expanded mineralization at 5 weeks after culture in osteoinductive medium (**Figure 5a**). Osteoblast mineralization, as a late phase of differentiation, was inhibited by TC and high concentrations (20 μg/ml) of OC, DC and MC (**Figure 5b**). Osteoblast mineralization was promoted by low (5 μg/ml) and moderate (10 μg/ml) concentrations of OC, DC and MC.

**Figure 5.**
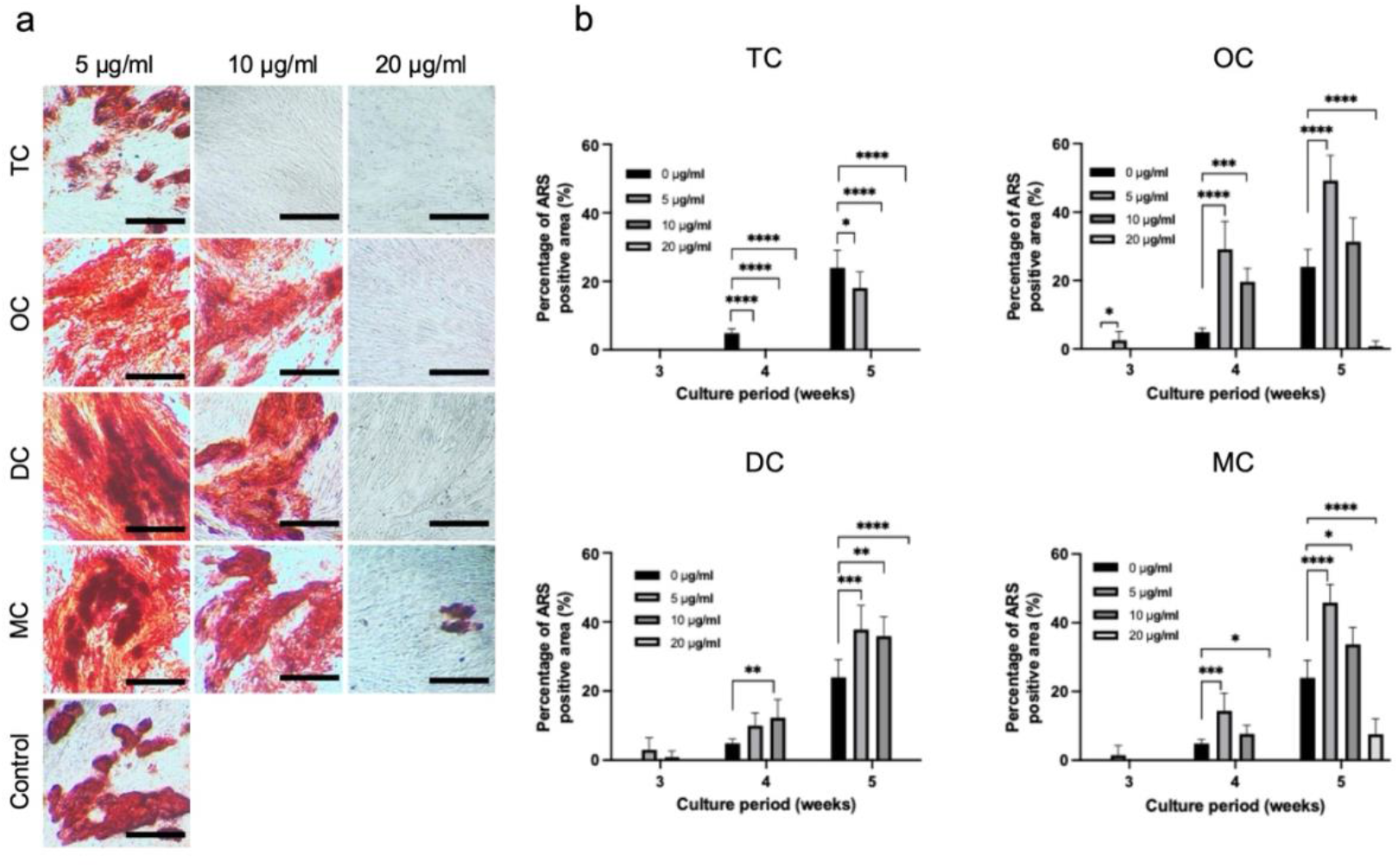
Effects of TC derivatives on osteoblast mineralization during differentiation. (a) Representative images of osteoblast cultures in osteoinductive medium showing the progression of mineralization in control cells. Initial mineral deposition was observed at 4 weeks, followed by extensive mineralized nodule formation at 5 weeks. (b) Mineralization of osteoblasts cultured with different TC derivatives at varying concentrations. TC and high concentrations (20 μg/mL) of OC, DC, and MC inhibited mineralized nodule formation, whereas low (5 μg/mL) and moderate (10 μg/mL) concentrations of OC, DC, and MC enhanced mineralization, indicating concentration-dependent regulation of late-stage osteoblast differentiation. Data are shown as the mean ± SD (n=6). \**p* < 0.02, \*\**p* < 0.005, *** *p* < 0.002, \*\*\*\**p* < 0.0001 compared with the control.

The cellular localization of various TC derivatives in bone nodules produced by osteoblasts was examined using high-resolution confocal laser microscopy (**Figure 6**). Fluorescence imaging revealed an extensive distribution of nuclei, visualized in blue by Hoechst staining, across the entire surface of the glass coverslips. This uniform presence of nuclei indicated that the osteoblasts had reached confluence, thereby establishing a suitable environment for bone nodule formation and subsequent mineral deposition. The merged images demonstrated that all the tested TC derivatives were distributed in the mineralized regions of the bone nodules. These mineral deposits were identified by positive staining with ARS, confirming the presence of mineralized extracellular matrix. Furthermore, in the corresponding bright-field images acquired without a fluorescence filter, mineralized regions appeared as distinct, needle-like black structures, providing complementary morphological evidence of mineral deposition. The colocalization of TC derivatives with ARS-positive areas suggests that TC derivatives preferentially accumulate in mineralized bone matrix produced by osteoblasts.

**Figure 6.**
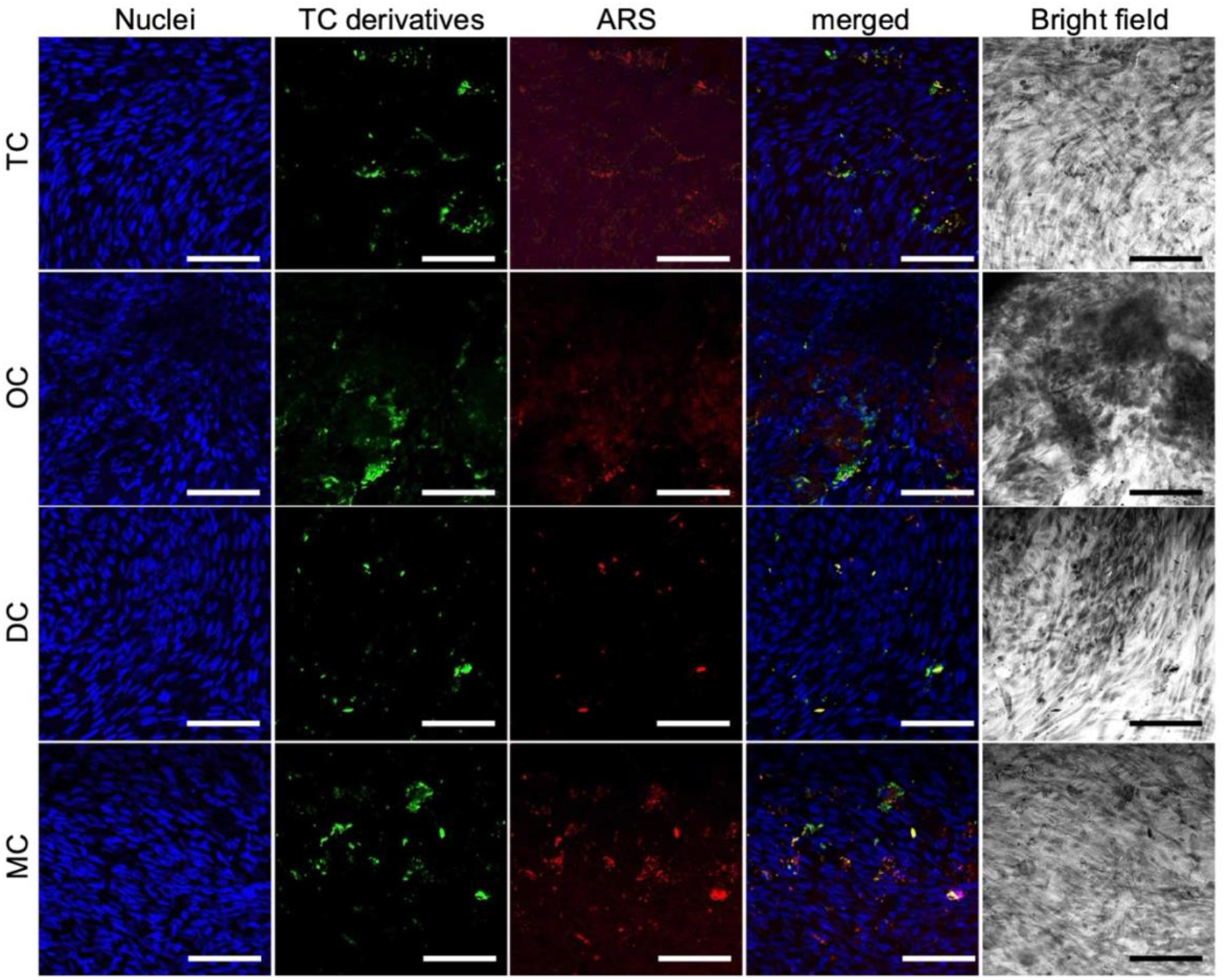
Localization of TC derivatives within mineralized bone nodules formed by osteoblasts. High-resolution confocal laser microscopy images showing the distribution of TC derivatives in osteoblast cultures. Cell nuclei were stained with Hoechst (blue), indicating a confluent cell layer across the coverslips. Mineralized extracellular matrix was identified by ARS staining (red). Merged fluorescence images revealed co-localization of TC derivative signals with ARS-positive mineralized regions, demonstrating preferential accumulation of TC derivatives within bone nodules. Corresponding bright-field images showed mineralized areas as needle-like black structures, providing morphological confirmation of mineral deposition. Fluorescence images of osteoblasts cultured with 10 µg/ml TC, OC, DC, and MC for 5 weeks in osteoinductive medium as visualized by nuclei (blue), TC derivatives (green) and mineral deposits stained with ARS (red). Scale bar=10 *μ*m.

### 2.3. Zebrafish as a model of bone development

To compare the effects of TC derivatives on bone development, a zebrafish model was used to expose embryos to the different types of TC derivatives at several concentrations (Figure 7). Fluorescence imaging was performed to assess skeletal development, with a particular focus on the cleithrum and operculum regions, which are key indicators of early craniofacial bone formation. At 4 dpf, both bone structures were clearly developed and visualized under fluorescence microscopy in control embryos, confirming normal skeletal growth. In contrast, embryos treated with TC derivatives exhibited inhibited bone development in a concentration-dependent manner (**Figure 7a**), with TC presented as a representative example.

**Figure 7.**
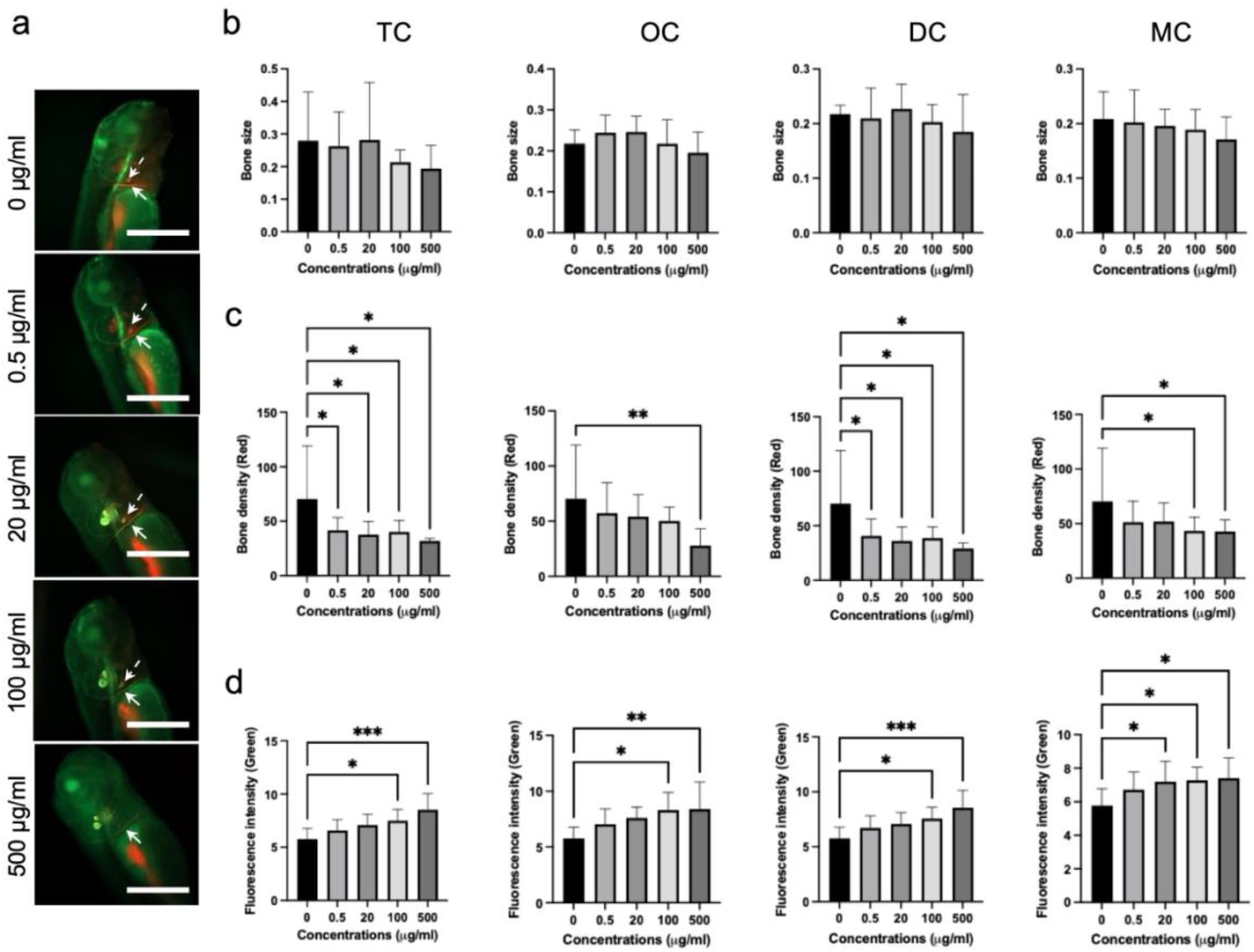
Effects of TC derivatives on craniofacial bone development in zebrafish embryos. (a) Representative fluorescence images of zebrafish embryos at 4 days post-fertilization (dpf) showing the cleithrum and operculum following exposure to increasing concentrations of TC derivatives (TC shown as a representative example). Mineralized bone structures were visualized by fluorescence staining. The cleithrum is marked with arrows, and the operculum is marked with dotted-line arrows. Scale bar=100 *μ*m. (b) Quantification of cleithrum area in embryos treated with different concentrations of TC derivatives, showing no significant changes in bone size. (c) Quantitative analysis of mineralized bone density within the cleithrum region (red fluorescence channel), demonstrating a significant dose-dependent decrease with increasing TC derivative concentrations. (d) Quantification of TC derivative accumulation within the cleithrum region (green fluorescence channel), showing a significant dose-dependent increase in fluorescence intensity, indicating progressive compound localization in developing bone tissue. Data are shown as the mean ± SD (n=8-18). \**p* < 0.02, \*\**p* < 0.005, *** *p* < 0.002 compared with the control.

Quantitative analysis revealed that the overall size of the cleithrum did not show significant differences across the tested concentrations of TC derivatives (**Figure 7b**). However, a marked decrease in cleithrum bone density was observed with increasing concentrations of the derivatives (**Figure 7c**), suggesting impaired mineralization during bone formation. Conversely, fluorescence intensity associated with TC derivatives within the cleithrum region increased significantly in a dose-dependent manner (**Figure 7d**), indicating progressive accumulation of these compounds in developing bone tissue. Collectively, these results demonstrate that TC derivatives preferentially localize to mineralized structures while simultaneously disrupting normal craniofacial bone mineralization in zebrafish embryos.

## 3. Discussion

### 3.1. Effects on osteoclastogenesis

TCs and their derivatives are known to induce apoptosis in rabbit osteoclasts and inhibit bone resorption by osteoclasts ^[10,19]^. In this study, all tested TC derivatives inhibited osteoclast differentiation and bone resorption, as indicated by TRAP staining and measurements of resorption pits (**Figures 1 and 2**). Dose-dependent inhibition was observed in osteoclasts cultured with TC, OC, and MC. DC inhibited osteoclast resorption at low, moderate and high concentrations.

Osteoclasts are formed by the fusion of monocytic precursors with stimulation by M-CSF and RANKL and then activated to resorb bone by secreting acid to dissolve the inorganic components of the bone matrix (hydroxyapatite minerals) and proteinase to degrade the organic components of the bone matrix, such as cathepsin K and MMPs ^[20,21]^. Osteoclasts produce various types of MMPs to degrade bone ECM, including MMP-1, MMP-2, MMP-3, MMP-9, MMP-10, MMP-12, MMP-13, and MMP-14 ^[21,22]^. MMP-2 has been reported to degrade type I collagen in the bone matrix, facilitate osteoclastogenesis and amplify various signaling pathways that enhance osteolysis in bone metastasis ^[23–25]^. MMP-9 is among the important proteinases that degrade the organic matrix and is secreted into resorption lacunae by osteoclasts; additionally, MMP-9 participates in osteoclast recruitment and the release of growth factors from bone ECM ^[21,24,25]^. In mouse RAW264.7 cells, osteoclast differentiation is inhibited via the inhibition of MMP-9 by DC (2 μg/ml) ^[26]^.

The results of the present study reveal that the secretion of MMP-2 and MMP-9 by human osteoclasts was inhibited by the addition of TC and derivatives such as OC, DC, and MC (**Figure 3**). These results indicate that these compounds inhibit osteoclastogenesis through the inhibition of MMP-2 and MMP-9. Analysis of resorption pit morphology on bone slices revealed that all tested compounds reduced both the volume and depth of bone resorption. TC, OC, and MC displayed dose-dependent inhibition, whereas DC demonstrated a potent effect even at the lowest concentration. This finding is consistent with the literature, which highlights the ability of DC to inhibit MMP-9 and MMP-2, both of which are critical enzymes in osteoclast-mediated matrix degradation ^[23–25]^. These enzymes degrade type I collagen and other matrix components, facilitating bone resorption. The observed downregulation of MMP secretion—particularly the strong suppression of MMP-9 across all compounds and doses— corroborates this mechanism and highlights a shared inhibitory pathway among tetracyclines.

These findings support prior reports indicating that TC derivatives can suppress osteoclastogenesis, potentially by inhibiting nuclear factor-kappa B (NF-κB) signaling ^[11]^ and MMP activity ^[26,11]^. Tissue inhibitors of metalloproteinases (TIMPs) play crucial roles in regulating the activity of MMPs in osteoclasts ^[27]^. The balance between TIMPs and MMPs is essential for proper bone remodeling. Disruption of this balance, either through excessive MMP activity or insufficient TIMP activity, can lead to various pathological conditions, such as rheumatoid arthritis, cancer, and periodontitis ^[28,29]^. It would be beneficial to study the expression of TIMPs to consider drug design for proper bone remodeling in future studies.

### 3.2. Effects on osteoblast proliferation and differentiation

The results of the present study reveal that the tested compounds did not negatively affect initial cell adhesion (**Figure 4**). Osteoblast proliferation assays revealed that low and moderate concentrations of TC, DC, and MC increased proliferation at early stages (3 d), whereas only TC had a significant effect at 7 d. High concentrations of tetracyclines (60–80 mg/ml) can inhibit human osteoblast proliferation because of the destruction of mitochondria ^[30]^. These results indicate that there is a suitable concentration to exert positive effects on osteoblasts.

The effect on osteoblast differentiation, as measured by mineralized matrix formation (**Figure 5**), was concentration dependent and drug specific. The high concentration (20 μg/ml) of all the tested tetracyclines inhibited mineralization, whereas the low and moderate doses of OC, DC, and MC promoted mineralization. Interestingly, compared with the other semisynthetic derivatives, TC appeared to suppress mineralization across all concentrations, suggesting a negative effect. TC derivatives promote osteogenic differentiation through different pathways in human mesenchymal stem cells; TC and DC activate the Wnt pathway, whereas MC upregulates Hedgehog signaling ^[31]^. TC and DC also stimulate actin remodeling processes, inducing morphological changes in osteogenic differentiation and the expression of alkaline phosphatase, RUNX2, SP7, and SPARC as differentiation markers ^[31]^. Bisphosphonates are the most common drugs for osteoporosis treatment and inhibit various types of MMPs (MMP-1, MMP-2, MMP-3, MMP-7, MMP-8, MMP-9, MMP-12, MMP-13, and MMP-14) in human osteoblast-like MG-63 cells ^[32]^. Thus, TC derivatives have potential as candidates for targeted therapy in bone-related disorders such as rheumatoid arthritis ^[33]^ and for preventing aseptic loosening of prosthetic joint implants ^[34]^.

Fluorescence imaging by confocal laser microscopy confirmed that the TC derivatives were localized within mineral deposits (**Figure 6**), further supporting their active integration into bone tissue and potential for long-term effects on bone remodeling. These results not only demonstrate the bone-targeting ability of TC derivatives at the cellular level but also highlight their potential utility as molecular probes or delivery vehicles for therapeutic agents aimed at mineralized tissues ^[35]^. This property may prolong the therapeutic duration but also necessitates cautious dose regulation to avoid long-term suppression of bone turnover.

### 3.3. Zebrafish as a model for bone development

Zebrafish embryos provide a valuable in vivo model for studying bone development because of their optical transparency, rapid growth, and conserved genetic pathways in skeletal biology ^[36]^. In this study, exposure to TC derivatives during early embryonic development led to dose-dependent inhibition of cleithrum bone formation (**Figure 7**). This effect was most prominent at higher concentrations (100–500 μg/ml), suggesting the importance of dose in achieving a therapeutic effect without impairing developmental processes. The inclusion of zebrafish data strengthens the translational relevance of our findings and highlights potential risks in bone development.

The present results suggest that TC derivatives, particularly DC and MC, may be repurposed as dual-action therapeutic agents for osteoporosis capable of inhibiting osteoclast-mediated bone resorption and promoting osteoblast function at appropriate doses. The observed biphasic effect aligns with the notion that many pharmacological agents exhibit dose-dependent dual effects: stimulatory effects at low concentrations and inhibitory effects at high doses. These findings indicate that if repurposed for osteoporosis, TC derivatives must be used at carefully optimized dosages to maximize osteoanabolic effects while minimizing inhibitory effects.

Nonetheless, several important limitations must be addressed before clinical application. First, the in vitro and zebrafish models do not fully replicate the complexity of bone remodeling in mammalian systems. Future preclinical studies in rodent models of osteoporosis are necessary to confirm efficacy, biodistribution, and long-term safety. Additionally, potential systemic effects, such as immunosuppression, alterations to the gut microbiome, and antibiotic resistance, must be carefully evaluated even for chemically modified tetracyclines with reduced antimicrobial activity. Notably, the anti-collagenase activity of low dose doxycycline was clinically demonstrated and it is a part of periodontal treatment regimen in some countries, supporting the concept that TC derivatives can be repurposed for chronic, non-antibiotic applications. The design of nonantibiotic tetracycline analogs (e.g., chemically modified tetracyclines lacking the C4 dimethylamine group) offers a promising avenue to retain bone-modulating effects while eliminating antimicrobial activity. These analogs may allow chronic use without the drawbacks associated with conventional antibiotics.

## 4. Conclusion

This study demonstrates the strong potential of TC derivatives as modulators of bone remodeling. Their dual effects—suppressing osteoclastogenesis while promoting osteoblast proliferation and mineralization—position them as viable candidates for repurposing in osteoporosis therapy.

## 5. Materials and Methods

### 5.1. Osteoclast culture

The precursors of osteoclasts were obtained from human bone marrow. The donation protocol was approved by the Human Subjects Committee of the University of Turku. Monocytes were isolated from bone marrow from patients who underwent hip replacement surgery as previously described ^[27]^ (from 49 to 70 years old, female and male). Briefly, human bone marrow was obtained from the femoral collum and trochanteric region during the operation. The bone marrow samples were plated in 75-cm^2^ tissue culture flasks for 1 day. The nonadherent cells were used for osteoclast culture. After the nonadherent cells were collected, the flasks were washed with PBS once and then pooled into two cell fractions. Buffy coats layered over Ficoll-Paque solution were collected, washed with PBS twice and resuspended in α-minimum essential medium (α-MEM; Sigma‒Aldrich, MO, USA) supplemented with 10% fetal bovine serum (FBS; Gibco, MA, USA), L-glutamine (Gibco, MA, USA), and 100 U/ml penicillin–streptomycin (Gibco, MA, USA). The enriched mononuclear cells were seeded onto bovine bone slices at a density of 1×10^6^ cells/cm^2^ and cultured with 0, 5, 10, and 20 µg/ml TC (Sigma‒Aldrich, St. Louis, MO, USA), OC (Sigma‒Aldrich, St. Louis, MO, USA), DC (Sigma‒Aldrich, MO, USA), and MC (Sigma‒Aldrich, MO, USA). Osteoclasts were induced by the addition of 50 ng/mL RANKL (PeproTech, NJ, USA), 25 ng/mL macrophage colony-stimulating factor (M-CSF, R&D Systems, MN, USA), 5 ng/mL transforming growth factor beta (TGF-β, R&D Systems, MN, USA) and 1 μM dexamethasone (Sigma‒Aldrich, MO, USA) in a humidified atmosphere of 5% CO_2_ at 37°C for 14 days. Half of the medium was changed every 3–4 days.

### 5.2. MMP secretion from osteoclasts

Osteoclast precursors were cultured on bone slices with 0, 5, 10 and 20 µg/ml TC, OC, DC, and MC. After 14 days, the cells were lysed with 200 *μ*l of lysis buffer containing 50 mM Tris-HCl (pH 6), 150 mM NaCl and 1% Triton X–100. The cells were scraped, collected into tubes, and sonicated for 10 seconds. The cell lysates were frozen at –20 °C until use. The concentrations of MMP–9 and MMP–2 were detected with Luminex technique using commercial multiplex kits (Milliplex^®^ Human MMP Magnetic Bead Panel 2, HMMP2MAG 55K, Millipore Corporation, MA, USA).

### 5.3. Osteoclast differentiation and resorption

Osteoclast precursors were cultured on bovine bone slices for 14 days and fixed in 4% paraformaldehyde for 20 min. To confirm the differentiation of osteoclast precursors into osteoclasts, the cells were stained for tartrate-resistant acid phosphatase (TRAP, Sigma‒ Aldrich, MO, USA) and with Hoechst 33258 (Sigma‒Aldrich, MO, USA). The number of TRAP-positive multinuclear cells per unit area was counted on each bone slice. A total of at least 6 fields on 6 bone slices were counted to obtain an average for each experiment.

After each bone slice was subjected to fluorescence microscopy, the osteoclasts were delicately scrubbed and washed with distilled water. The resorption activity of the osteoclasts was assessed by measuring the morphological parameters of the resorption pits. The dried bone slices were examined with a color laser microscope (Olympus, OLS4100) equipped with an image analysis system. The depth and volume of resorption pits per unit area were measured to quantify the resorption efficacy of the osteoclasts. Measurements were performed for 5 areas on 6 slices to obtain an average for each experiment.

### 5.4. Osteoblast culture

Human osteoblast-like cells (MG-63) were maintained in Dulbecco’s modified eagle medium (DMEM; Gibco, MA, USA) supplemented with 10% FBS, 100 U/ml penicillin–streptomycin and 2mM L-glutamine in a humidified atmosphere of 5% CO_2_ at 37°C. After reaching 70% confluency, the cells were detached by the treatment with 0.05% trypsin-EDTA (Gibco MA, USA). The cell suspension was used for subculture or further experiments such as adhesion, migration, proliferation and differentiation assay. The medium was changed every 3–4 days.

### 5.5. Osteoblast proliferation

Cells were seeded into 96-well cell culture plates at a density of 5 × 10^3^ cells/cm^2^ and cultured with 0, 5, 10 and 20 µg/ml TC, OC, DC, and MC. After 1, 3 and 7 d, cell proliferation was evaluated using a WST-8 assay kit (Dojindo, CK04 34707621).

### 5.6. Osteoblast differentiation

Cells were seeded into 96-well cell culture plates at a density of 1×10^4^ cells/cm^2^ and cultured until confluency. The cells were cultured in osteoinductive medium supplemented with 100 nM dexamethasone (Sigma‒Aldrich, MO, USA), 10 mM β-glycerophosphate (Sigma‒Aldrich, MO, USA) and 50 µg/ml ascorbic acid (Sigma‒Aldrich, MO, USA) supplemented with 0, 5, 10 and 20 µg/ml TC, OC, DC, and MC in a humidified atmosphere of 5% CO_2_ at 37°C. After being cultured for 3, 4, and 5 weeks, the cells were fixed in 4% paraformaldehyde for 20 min. To visualize their differentiation into osteoblasts, the cells were stained with 40 mM alizarin red S (ARS, Sigma‒Aldrich, MO, USA) and observed under a fluorescence microscope (Olympus IX71). Whole images of each 96-well plate were captured to obtain the average percentage of ARS-positive areas and compare the effects of the TC derivatives on osteoblast differentiation.

### 5.7. Localization of TC derivatives in bone nodules

Cells were seeded onto glass coverslips in 48-well cell culture plates at a density of 5×10^3^ cells/cm^2^ and cultured until they reach confluence. The cells were cultured in osteoinductive medium supplemented with 100 nM dexamethasone, 10 mM β-glycerophosphate and 50 µg/ml ascorbic acid supplemented with 10 µg/ml TC, OC, DC, and MC in a humidified atmosphere of 5% CO_2_ at 37°C. After being cultured for 5 weeks, the cells were fixed in 4% paraformaldehyde for 20 min. To visualize the localization of TC derivatives in mineral deposits produced by osteoblasts, the cells were stained with Hoechst 33258 (Sigma‒Aldrich, MO, USA) and 40 mM ARS, mounted in glycerol and observed using a high-resolution confocal laser microscope (Zeiss LSM880) ^[37]^.

### 5.8. Zebrafish model

Wild-type (WT) zebrafish embryos were maintained at the Zebrafish Core Facility at Turku Bioscience. Several male and female adult zebrafish (1:1 ratios) were chosen and placed in individual breeding chambers for 24 h. At 0 days postfertilization (dpf), the adult fish were transferred back to home tank. The embryos were collected using a strainer and placed into a Petri dish with E3 medium containing 5 mM NaCl, 0.17 mM KCl, 0.33 mM CaCl_2_·2H_2_O, 0.33 mM MgCl_2_·6H_2_O (pH 7.2–7.3), and 0.57% methylene blue (Sigma‒Aldrich, MO, USA). The fertilized embryos were identified using a light microscope. The fertilized embryos were transferred into separate Petri dishes with E3 medium and 0.2 mM 1-phenyl 2-thiourea (PTU; Sigma‒Aldrich, MO, USA) to block melanin synthesis and render the embryos optically transparent. At 1 dpf, 2 mg/ml pronase (Sigma‒Aldrich, MO, USA) was added to the Petri dish to facilitate hatching. The Petri dish was kept in an incubator (+28.5 °C). The 2-dpf zebrafish embryos were exposed to 0, 0.5, 20, 100, and 500 µg/ml TC, OC, DC, and MC in E3 medium supplemented with 0.2 mM PTU for 2 days.

At 4 dpf, the zebrafish embryos were stained with 0.01% ARS (Sigma‒Aldrich, MO, USA) diluted in E3 medium (pH 7.4) ^[38]^ at 28.5 °C for 15 min. After the zebrafish embryos were washed twice with E3 medium, the embryos were kept in E3 medium supplemented with 0.2 mM PTU and 200 mg/l tricaine for anesthesia. The embryos were observed under a fluorescence microscope (Zeiss AxioZoom V16 Stereomicroscope). EGFP (green channel, excitation filter BP 470/40nm, emission filter BP 525/50nm) was used to visualize the TC derivatives, and Texas Red (red channel, excitation filter BP 560/40nm, emission filter BP 630/75nm) and Hamamatsu sCMOS Orca Flash4.0 LT + camera was used to visualize the ARS staining. The microscope was operated with ZEISS ZEN software (version 3.7), which was further used for image processing the raw image data. To quantitatively assess the effects of TC derivatives on zebrafish bone development, the cleithrum area as well as the fluorescence intensities corresponding to mineralized bone (red channel) and TC derivatives (green channel), were measured for each image using ImageJ software (version 1.54f).

### 5.9. Statistical analysis

Accurate quantification of the different samples was achieved by performing more than three independent experiments. Statistical analysis between groups was performed by analysis of variance (ANOVA) with the Tukey formula for post hoc multiple comparisons using the SPSS software package (version 29, Chicago, IL). A statistical significance level of p < 0.05 was used for all tests. All data are expressed as the mean ± standard deviation (SD).

## 6. Acknowledgements

This study was supported by the Sigrid Juselius Foundation, JSPS Grants-in-Aid for Scientific Research (JP23K08670), The Murata Foundation and the Turku Collegium for Science and Medicine. We thank Ms. Katja Sampalahti, Ms. Tatiana Peskova, and Dr. Vuokko Loimaranta for their technical assistance.

